# Biomimetic human lung alveolar interstitium chip with extended longevity

**DOI:** 10.1101/2022.12.23.521822

**Authors:** Kun Man, Jiafeng Liu, Cindy Liang, Christopher Corona, Michael D. Story, Brian Meckes, Yong Yang

## Abstract

Determining the mechanistic causes of lung diseases, developing new treatments thereof, and assessing toxicity whether from chemical exposures or engineered nanomaterials would benefit significantly from a preclinical human lung alveolar interstitium model of physiological relevance. The existing preclinical models have limitations because they fail to replicate the key anatomical and physiological characteristics of human alveoli. Thus, a human lung alveolar interstitium chip was developed to imitate key alveolar microenvironmental factors including: an electrospun nanofibrous membrane as the analogue of the basement membrane for co-culture of epithelial cells with fibroblasts embedded in 3D collagenous gels; physiologically relevant interstitial matrix stiffness; interstitial fluid flow; and 3D breathing-like mechanical stretch. The biomimetic chip substantially improved epithelial barrier function compared to transwell models. Moreover, the chip having a gel made of a collagen I-fibrin blend as the interstitial matrix sustained the interstitium integrity and further enhanced the epithelial barrier resulting in a longevity that extended beyond eight weeks. The assessment of multiwalled carbon nanotube toxicity on the chip was in line with the animal study.

## 1. Introduction

Being responsible for the exchange of oxygen and carbon dioxide, the alveoli, tiny air sacs in human lungs, also serve as the entry gateway for airborne particles such as engineered nanomaterials, air pollutants, and pathogens to enter the human body. They also provide a route for pulmonary drug delivery. As an example, engineered nanomaterials, especially when in their aerosolized form, can be inhaled into lungs potentially causing significant public health concerns [1]. It has been shown that inhaled carbon nanotubes (CNTs), a major class of engineered nanomaterials [2], cross the alveolar epithelial barrier and enter the interstitium, inducing progressive interstitial lung fibrosis in mice in weeks [3]. In addition to negative health implications, researchers seek to leverage the large alveolar surface area for inhalation-based delivery of drug-loaded nanoparticles, a cutting-edge technology for lung cancer therapy [4]. Thus, pathophysiologically relevant preclinical alveolus models are critical for understanding and treatment of lung diseases and toxicity assessment of engineered nanomaterials.

Current preclinical models (*e.g*., animal models, conventional 2D models, and lung organoids) have limitations. Differences in the anatomy, physiology, and genomics between animals and humans make conventional animal models inconsistent and inaccurate, limiting their translation to clinical studies [5]. In 2D models, human primary cells or cell lines are cultured on flat plastic surfaces, which leads to different behaviors compared to their *in vivo* counterparts that interferes with the replication of *in vivo* phenotypes or functions observed in native tissues [6]. Lung organoids replicate complex 3D structures and functions of the lung but cannot model some key *in vivo* lung microenvironmental features like the air-liquid interface (ALI) [7, 8]. Moreover, all these models lack physiologically relevant mechanical characteristics, such as interstitial flow and/or lung breathing movement.

Organ-on-a-chip systems can replicate organismal level function and have recently attracted great attention [9, 10]. The human lung chips model the ALI and alveolus-capillary interface, mimic breath movement [9], and have been adapted to model several diseases and to test therapeutic drugs [11, 12]. Several human lung fibrosis chips have also been developed to replicate stromal-vascular and stromal-epithelial interfaces [13, 14]. However, these chips do not mimic the basement membrane, interstitium stiffness, or breathing movement critical for lung functions and disease [15, 16]. Note that our recent study reveals that mechanical stretch dimensionality (*i.e*., 3D stretch similar to native alveoli) is critical to epithelium formation [17]. Due to the missing key alveolar microenvironmental features, the epithelial barrier function is not sustained, and most of these chips are only viable for 1-3 weeks [9, 11, 18, 19]; therefore, they are not suitable for investigation of chronic diseases.

In this study, we developed a human alveolar interstitium chip that consists of an electrospun nanofibrous membrane for co-culture of alveolar epithelial cells with lung fibroblasts encapsulated in 3D collagenous hydrogels. The chip also provides interstitial flow and 3D mechanical stretch of physiological relevance to the cells. The pore size of the nanofibrous membrane was optimized to promote epithelium formation. The interstitium chip exhibited enhanced epithelial barrier function with longevity extended beyond eight weeks. Furthermore, we assessed the penetration of multi-walled CNTs (MWCNTs) across the epithelium of the chips, which showed results consistent with the *in vivo* study [20].

## 2. Results and Discussion

### 2.1. Design and fabrication of the human alveolar interstitium chip

The functions of lung alveoli are profoundly affected by alveolar anatomical and physiological characteristics. Anatomically, the alveolar wall is a thin epithelium that faces the alveolar lumen and is covered with a thin fluid layer to form an ALI. Supported by a microporous basement membrane, the epithelium forms a barrier due to the presence of tight junctions and adherens junctions [21], which prevents access of nanomaterials to the subepithelium [22]. The interstitium surrounding the epithelium displays an interrelated framework of extracellular matrix (ECM) proteins such as collagen and elastin and is rich with interstitial fluids [23]. *In vivo* studies suggest that the penetration of nanomaterials into the lung interstitium and interaction with local fibroblasts may be a critical step in nanomaterial induced fibrogenesis [24]. A key cellular mechanism of fibrogenesis is fibroblast activation and subsequent induction of ECM (in particular, collagen) production and accumulation leading to fibrosis [25, 26]. Lung fibroblasts exhibit the stiffness-dependent fibrogenic responses to MWCNTs [27]. In addition, interstitial fluid flow could alter cytoskeletal organization, influence cell proliferation, and remodel the interstitial matrix [28, 29]. Moreover, the breathing movements expose the epithelium and interstitium to cyclic 3D stretching and relaxation. The mechanical stretch is known to have a profound influence on the formation and function of cells, tissues and organs [30].

To establish the primary functions of lung alveoli, we developed a dynamic human lung alveolar interstitium-on-a-chip (interstitium chip) system which closely replicates the key anatomical (nanofibrous membrane to imitate the basement membrane) and physiological (interstitial matrix stiffness, interstitial flow, and cyclic 3D breathing-like mechanical stretch) characteristics of human lung alveoli (Fig. 1).

**Fig. 1.**
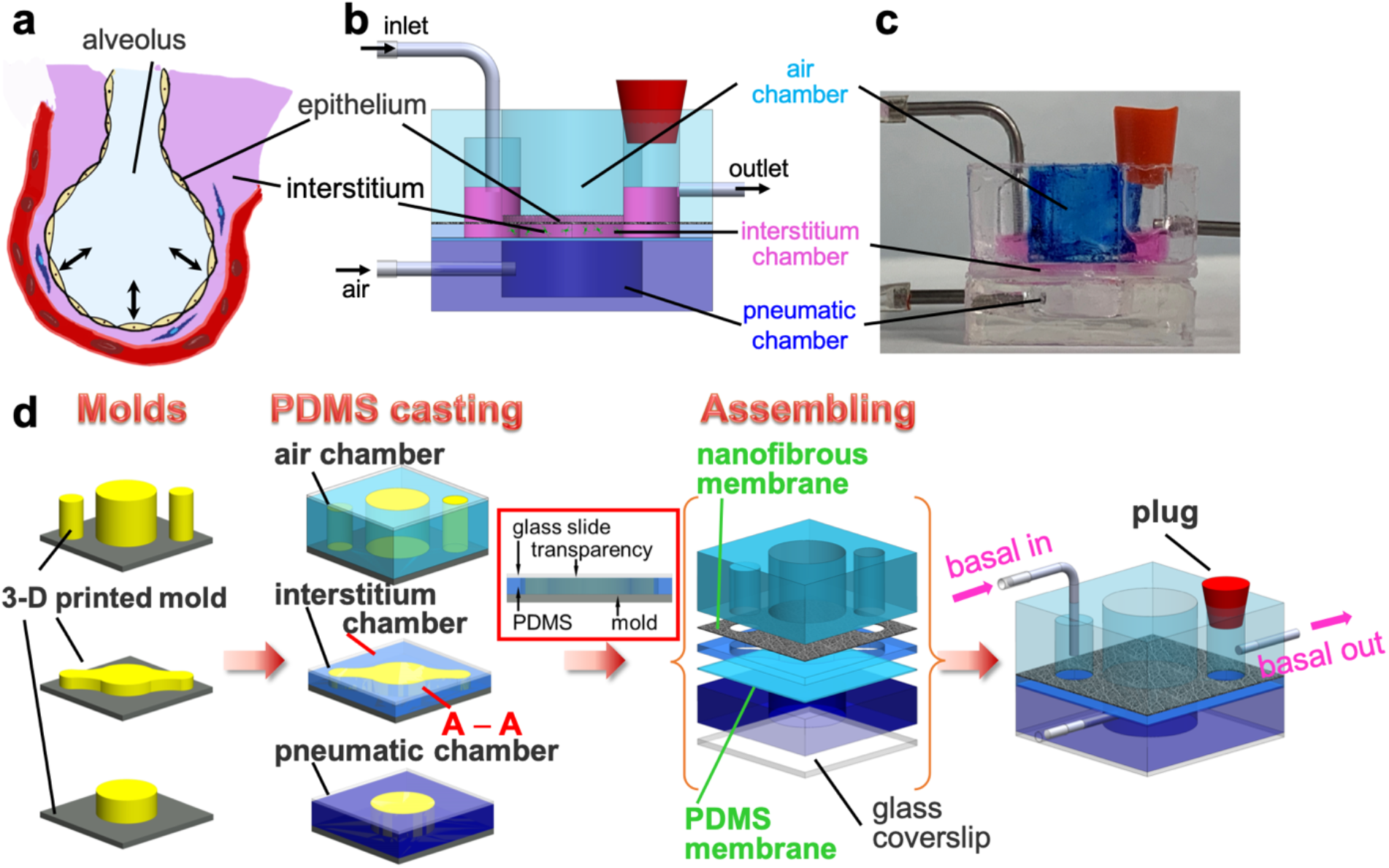
Design and fabrication of human lung alveolar interstitium chip. (a) Schematic illustration of human lung alveolus. (b) Design, (c) photo, and (d) fabrication of the chip. The red box in (d) is the cross-sectional view of A-A of the interstitium chamber fabrication.

The chip consisted of a co-culture section and a pneumatic section, both made of polydimethylsiloxane (PDMS). The co-culture section was comprised of an air (apical) chamber and an interstitium (basal) chamber, between which an electrospun nanofibrous membrane (~10 μm thick) was sandwiched (Fig. 1d). Human lung alveolar epithelial cells formed a tight epithelium on the nanofibrous membrane at the ALI in the air chamber. Human lung fibroblasts were embedded in a 3D collagenous matrix in the interstitium chamber. The pneumatic section was formed by permanently bonding a PDMS membrane (15 μm thick) on the pneumatic chamber.

The stiffness of the interstitial matrix was primarily a consequence of using a type I collagen (Col I) gel for which the stiffness of the gel can be controlled, with values ranging between 1 kPa and 100 kPa, by adjusting the concentration of Col I solution between 3 and 60 mg/mL, respectively [31]. This range of stiffness covers normal (1-5 kPa) and fibrotic (20-100 kPa) human lung tissues [32–34]. A syringe pump was used to perfuse culture media continuously into the interstitium chamber to generate an interstitial flow of physiological relevance [35–38], as demonstrated in our computational simulation (see Supporting Materials, Fig. S1).

Our previous study revealed that physiologically relevant cyclic 3D mechanical stretch significantly upregulated the expression of tight and adhesion junction proteins compared to 1D and 2D stretches and static controls [17]. A circular PDMS membrane was thus designed for the pneumatic chamber to provide 3D stretch. When air was pumped in and withdrawn from the pneumatic chamber, the PDMS membrane, the interstitium, and the nanofibrous membrane moved up and down, generating 3D radial stretch to mimic breathing movements. There was no significant difference in the strain between the experimental observation and the theoretical calculation (Fig. S2, Video S1 and S2). Thus, the mechanical stretch was transmitted through the interstitium to the nanofibrous membrane, which exposed the fibroblasts in the interstitium and the epithelial cells on the nanofibrous membrane to the designated strain. The level of applied strain ranges from 5% to 15% at a frequency of 0.2 - 0.3 Hz (12-20 time/min) to match normal levels of strain observed in alveoli within the whole human lung *in vivo* [39].

These anatomical (nanofibrous membrane) and physiological (interstitium matrix stiffness, interstitial flow, and cyclic 3D stretch) characteristics of the interstitium chip could be adjusted and optimized to promote and sustain chip function. In addition, the air chamber, electrospun nanofibrous membrane, and interstitium chamber were assembled into the co-culture section by applying the microtransfer assembly (μTA) technique that we previously developed based on microcontact printing with PDMS prepolymer thin film as the adhesive [40]. This technique allowed us to adjust the adhesive strength between the chip sections by controlling the extent of curing of the PDMS thin film which then allowed for the selective separation of sections for downstream analysis.

### 2.2. Nanofibrous membrane promoted the formation of a tight epithelium

Nanofibrous membranes of various pore sizes were generated by electrospinning polycaprolactone (PCL) solution. By controlling the electrospinning time, nanofibrous membranes of three different pore sizes were fabricated, *i.e*., small pores (S, 2.32 ± 1.52 μm^2^), medium pores (M, 8.75 ± 5.79 μm^2^), and large pores (L, 15.63 ± 9.79 μm^2^) (Fig. 2). Because all other processing parameters were kept unchanged, the fiber diameters of all of these membranes were similar, *i.e*., 890 ± 230 nm, 880 ± 240 nm, and 880 ± 210 nm for the small, medium, and large pore size membranes, respectively. Therefore, these membranes allowed us to investigate the effects of pore size on epithelium formation.

**Fig. 2.**
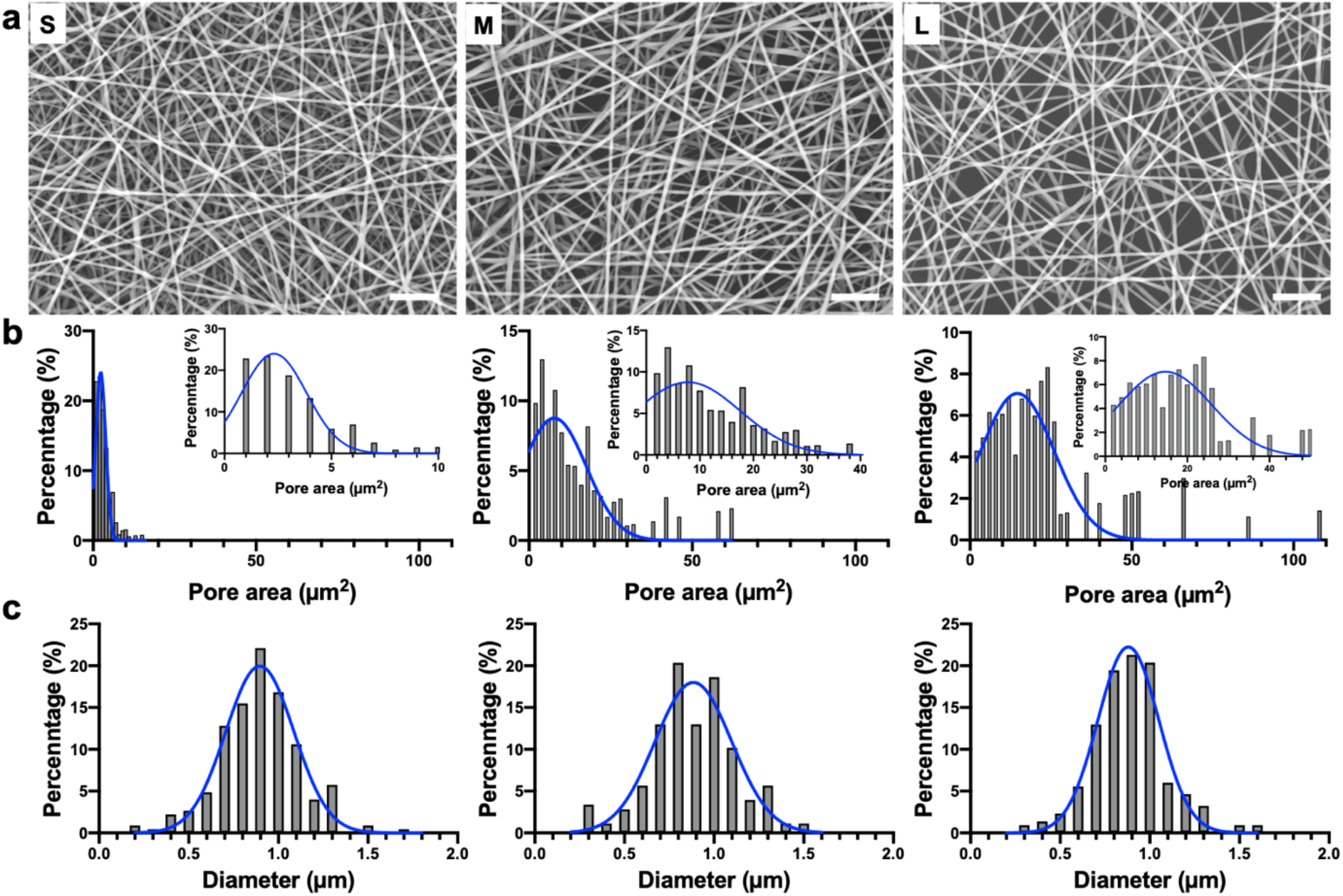
Characterization of electrospun nanofibrous membranes. (a) Scanning electron micrographs (SEM) of nanofibrous membranes of small (S), medium (M), and large (L) pore sizes. Scale bars: 20 μm. (b) Pore size distributions of these nanofibrous membranes. The insets magnify the main pore size distributions. (c) Fiber diameter distributions of these nanofibrous membranes.

Human alveolar epithelial A549 cells were grown on these nanofibrous membranes while using a PCL flat surface as the control. After 10 days, the epithelial cells displayed continuous tight junction protein zonula occludens (ZO)-1 on the nanofibrous membranes compared to discrete ZO-1 expression on the flat control (Fig. 3a). The pore size effects on the expression of tight junction proteins ZO-1, ZO-3, and occludin and adhesion junction protein E-cadherin were evaluated by western blot. In general, nanofibrous membranes upregulated protein expression and the cells showed significantly higher expressions of ZO-1 and occludin on the medium and large pore membranes compared to the flat control (Fig. 3b).

**Fig. 3.**
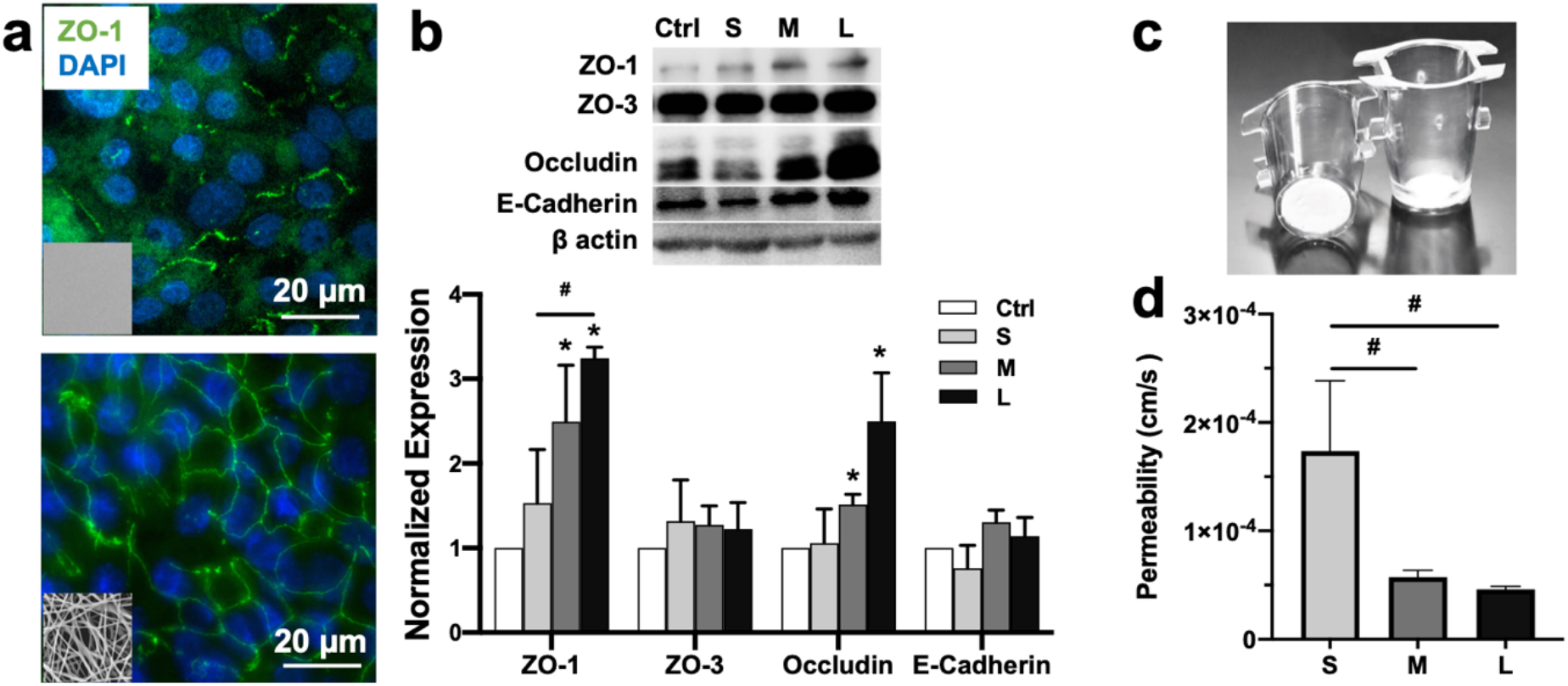
Nanofibrous membrane promoted the formation of tight epithelium. (a) Representative immunofluorescence images of A549 cells cultured on the PCL flat control and nanofibrous membranes (SEM images are presented in the insets) for 10 days. Green: ZO-1; blue: nuclei. (b) Western blotting of ZO-1, ZO-3, occludin, and E-cadherin of A549 cells cultured on the flat control and nanofibrous membranes with small (S), medium (M), and large (L) pore sizes (n = 3). The data were normalized to the mean value of the flat control. (c) Photos of transwell inserts attached with the nanofibrous membrane. (d) Permeability of 4kDa dextran across the epithelial cell layer on the nanofibrous membranes. ^*^: *p* < 0.05 compared to the flat control. ^#^: *p* < 0.05 between groups.

In addition, the nanofibrous membranes were attached to the transwell insert and the permeability coefficient of dextran – commonly used to assess transport via the paracellular route through tight junctions [41] across the epithelial monolayers was assessed. Similarly, the epithelium permeability of the nanofibrous membranes of medium and large sizes were significantly lower than that of the small pore membrane (Fig. 3d).

The basement membrane of lung alveolar epithelium exhibits oval-shaped pores of 0.75 to 3.85 μm in size and the microporous membrane facilitates type II pneumocytes to form a monolayer, enable heterocellular crosstalk, and act as a route for immune cells to move between the epithelium and interstitium [21, 42, 43]. PCL fibrous membranes have been shown to promote the barrier function of lung alveolar epithelium [44]. However, it was unclear if the pore size of fibrous membranes influenced alveolar epithelium barrier function. In this study, PCL nanofibrous membranes with various pore sizes were fabricated and their promoting effects on epithelium formation and permeability were verified. Occludin is linked via zonula occludens protein complexes to tight junctions that seal neighboring epithelial cells and limit paracellular diffusion. The upregulated expression of occludin and ZO-1 proteins and enhanced epithelial permeability on the medium and large pore membranes suggested that both nanofibrous membranes promoted the formation of tight alveolar epithelium. Although the large pore membrane showed a better promoting effect on tight epithelium, there was occasional leakage due to some large pores. Thus, the nanofibrous membrane of medium pores (8.75 ± 5.79 μm^2^) was selected for the following studies.

### 2.3. Alveolar environmental factors enhanced the epithelial barrier function

Next, we constructed the interstitium chip with the nanofibrous membrane of medium pore size, constrained by the other key physiological parameters, namely, the interstitium matrix stiffness, interstitial fluid, and cyclic 3D mechanical strain. Collagen is the most abundant protein in the interstitium matrix and thus Col I gel was used as the interstitial matrix for 3D culturing of the normal human lung fibroblasts (NHLFs) on the chip. Our previous study highlighted the significance of matrix stiffness in the fibrogenic responses of lung fibroblasts to MWCNTs [27]. On the chip the concentration of Col I was adjusted to 3 mg/mL to form a gel of 1 kPa in stiffness [45] to resemble the stiffness of normal lung tissue [33]. The interstitial medium perfusion rate was set as 20 μl/hr, which generated an average interstitial fluidic velocity of 0.6 – 1.4 μm/s (also see Fig. S1c-f), within the physiological range, *i.e*., 0.1 – 4.0 μm/s [37, 38]. To replicate the cyclic strain that lung alveoli experience during breath movement, which is 5 – 15% linear strain at a frequency of 0.2 – 0.3 Hz in normal conditions, a theoretical maximum 15% linear strain was applied at a frequency of 0.2 Hz. For comparison, the conventional transwell model was built as the controls. In the transwell model, the epithelial and fibroblast cells were grown on the apical and basal side of the insert membrane, respectively.

After one week of culture, the epithelial cells formed a dense epithelial layer on the chip; however, the gaps between the epithelial cells were evident in the transwell model (Fig. 4a, b). The epithelium permeability was evaluated using dextrans of 4 kDa and 70 kDa in molecular weight, which represents the intermediate and large-size agents transported through tight junctions [41, 46]. As shown in Fig. 4c, the permeability of both dextrans through cells grown on the chip was about 3~4 fold lower than that of the transwell model, *i.e*., 2.16±0.51 × 10^-5^ vs. 6.12±2.58 × 10^-5^ cm/s for 4 kDa dextran, and 7.79±1.75 × 10^-6^ vs. 2.97±1.21 × 10^-5^ cm/s for 70 kDa dextran. Note that the epithelium permeability of 4 kDa dextran on the chip was also improved by 2-fold compared to the monoculture on the nanofibrous membrane, *i.e*., 2.16 ± 0.51 × 10^-5^ vs. 5.73±2.70 × 10-^5^ (Fig. 3d, Table 1).

**Fig. 4.**
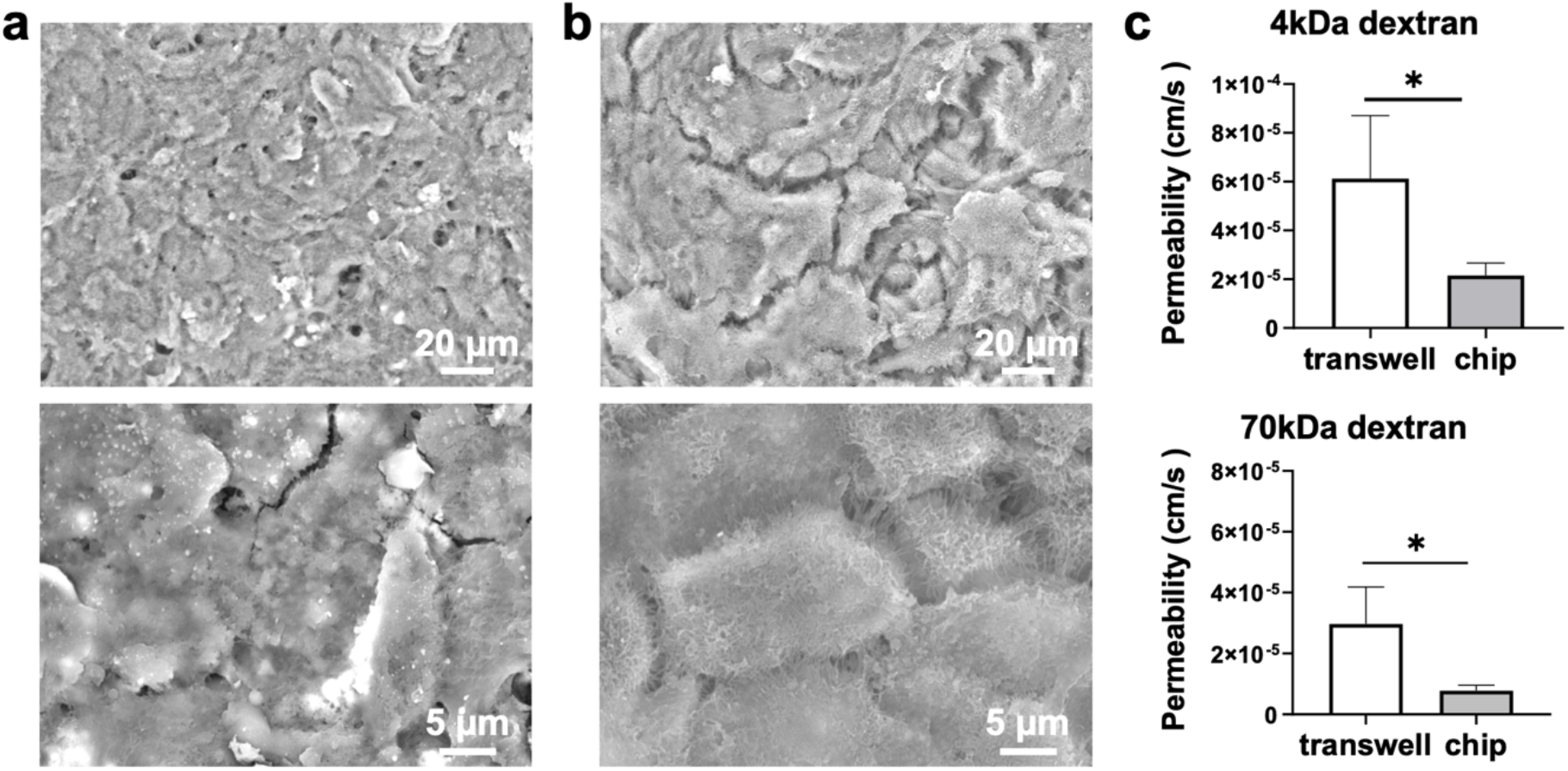
Interstitium chip promoted epithelial barrier function. Low and high magnification SEM images of the epithelial monolayer (a) on the chip and (b) in the transwell model. (c) Comparison of dextran permeability on the chip and the transwell model after 1-week of culture. ^*^: *p* < 0.05 compared to transwell model.

**Table 1.** Permeability summary

| Model | Duration | Permeability (cm/s) |  |
| --- | --- | --- | --- |
|  |  | 4 kDa dextran | 70 kDa dextran |
| Interstitium chip | 4 weeks | $7.08 \pm 1.24 \times 10^{-6}$ | $9.77 \pm 0.84 \times 10^{-8}$ |
| | 1 week | $2.16 \pm 0.51 \times 10^{-5}$ | $7.79 \pm 1.75 \times 10^{-6}$ |
| Transwell co-culture | 4 weeks | $4.96 \pm 3.39 \times 10^{-5}$ | $1.73 \pm 0.40 \times 10^{-6}$ |
| | 1 week | $6.12 \pm 2.58 \times 10^{-5}$ | $2.97 \pm 1.21 \times 10^{-5}$ |
| Nanofibrous membrane only | 1 week | $5.73 \pm 2.73 \times 10^{-5}$ | - |
| Literature reports [41, 47, 48] | | $10^{-6} - 10^{-5}$ | $10^{-7} - 10^{-6}$ |

In brief, the biomimetic interstitium chip demonstrated enhanced epithelial barrier function compared to a conventional transwell model and monoculture on the nanofibrous membrane. This enhancement was likely attributed to the synergy of the nanofibrous membrane, co-culture of the epithelial and fibroblast cells in a 3D gel, and physiologically relevant interstitium matrix stiffness, interstitial flow, and cyclic 3-D stretch.

### 2.4. The Col I-fibrin interstitial matrix sustained the chip

After two weeks of culture, interstitial matrix degradation and remodeling was observed on the chip. It is known that collagenous gels degrade relatively fast, and fibroblasts caused collagen gel contraction [49]. To alleviate the interstitium deterioration, fibrin was utilized to reinforce the collagen gel because fibrin gels have been reported to be stable for at least 12 months [50]. The degradation of fibrin gels, a process called fibrinolysis, can be controlled by inhibiters such as ε-aminocaproic acid (EACA) [51]. Additionally, fibrin gels can facilitate cell adhesion and promote fibroblasts to synthesize more collagen and other ECM proteins compared to collagen gels [52, 53]. Compared with pure collagen and fibrin gels, the Col I-fibrin blend gels exhibit intermediate properties [54]. Therefore, we prepared Col I-fibrin blend gels by mixing Col I solution and fibrin solution at volume ratios from 1:0.1 to 1:1 for 3D culturing of the lung fibroblasts. After culture under static conditions for two weeks, no gel deformation was observed in all blend gels, the fibroblasts spread, and cell elongation increased with an increase in the fibrin concentration (Fig. S3). The observation indicated that adding fibrin improved the interstitium integrity and likely extended chip longevity. To maintain matrix stability while maximizing collagen content, a Col I-fibrin gel with a ratio of 1:0.3 was used for long term culture of the chip.

The epithelial cells formed a monolayer after one week of culture on the chip with Col I-fibrin gel as the interstitial matrix. The epithelial cell growth medium was then aspirated from the air chamber to form an ALI and a mixed epithelial and fibroblast culture media (1:1, v/v) was perfused into the interstitium chamber. Three weeks later, a tight epithelium formed and the A549 cells displayed cuboidal shape and presented apical microvilli – the typical ultrastructural feature of human alveolar epithelial type II (ATII) cells (Fig. 5a, b). The layered structure of epithelium, secreted ECM, and electrospun nanofibrous membrane was evident in Fig. 5c. Importantly, the epithelium permeability was further enhanced. For example, for 4 kDa dextran, the permeability decreased by 3-fold from 2.16±0.51 × 10^-5^ cm/s (1 week) to 7.08±1.24 × 10^-6^ cm/s (4 weeks), significantly lower than 4.96±3.39 × 10^-5^ cm/s for the transwell model. For 70 kDa dextran, the permeability decreased by about 80-fold, from 7.79±1.75 × 10^-6^ cm/s (1 week) to 9.77±0.84 × 10^-8^ cm/s (4 weeks) compared to 1.73±0.40 × 10^-6^ cm/s for the transwell model (Table 1, Fig. 5d). The enhanced epithelium permeability values were comparable or lower than literature reports using transwell models and microfluidic devices [41, 47, 48].

**Fig. 5.**
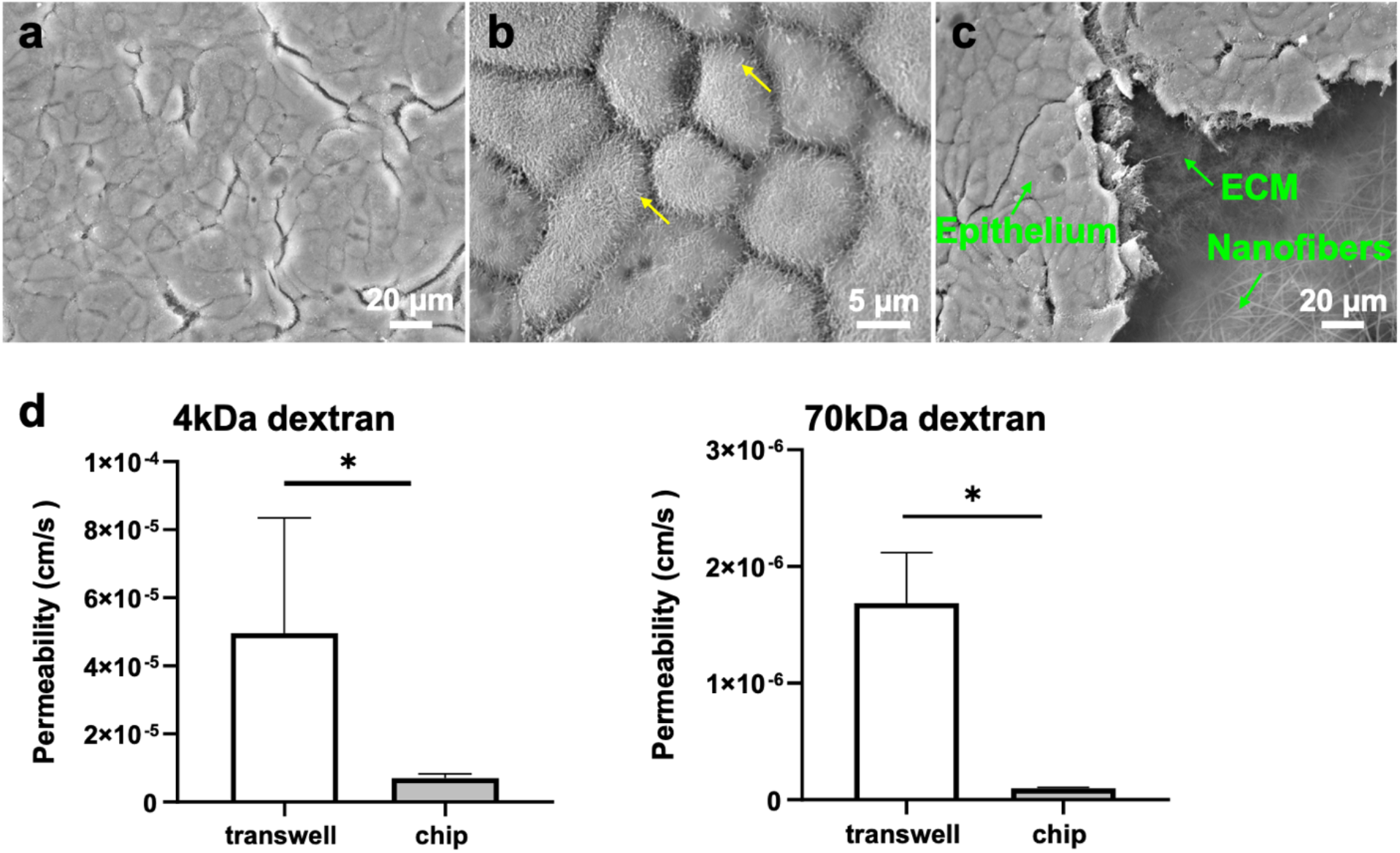
Long term culture of the lung interstitium chip. (a) Low and (b) high magnification SEM images of the epithelium on the chip. The yellow arrows in (b) point to the apical microvilli. (c) SEM image of the layered structure of the epithelium on the chip. (d) Dextran permeability of the epithelium after 4-week culture in the chip. ^*^: *p* < 0.05 compared to the transwell models.

A549 cells feature some of the ATII cell properties and are often used in *in vitro* lung alveolar models [47]. It was reported that ATII cells cocultured in transwells with lung fibroblasts embedded in a 3D matrix maintained their distinct phenotype for 7 days; conversely, the cells lost their phenotype within 3-5 days on conventional 2D models [55, 56]. On this interstitium chip, the A549 cells formed a tight epithelium in one week and displayed distinct ATII cell structural features like cuboidal shape and apical microvilli. Moreover, the epithelium permeability was significantly lower than the transwell models. Of note, no evident deterioration of the chip was observed after 8-weeks of culture; conversely, individual epithelial cells instead of a continuous cell monolayer was observed in the transwell model (Fig. S4).

The biomimetic interstitium chip demonstrated an enhanced epithelial barrier function with extended longevity. As fibrin is known to be formed by enzymatic cleavage and reorganization of fibrinogen and thrombin [57], interstitium integrity can be enhanced by reducing gel degradation via the administration of EACA, which blocks the binding of plasmin or plasminogen to fibrin [51].

A549 cells provide a reliable and straightforward cell source for chip development. However, the cells are from a lung tumor, are likely genetically unstable, and might not fully recapitulate epithelial barrier function *in vivo*. The use of primary human alveolar epithelial cells or epithelial basal stem cells will facilitate the establishment of a more *in vivo* like model [58]. Human macrophages can also be co-cultured with the epithelial cells at the ALI to facilitate the maintenance of lung homeostasis [59]. Taken together, the key alveolar microenvironmental factors can be fine-tuned to further improve chip longevity and performance.

### 2.5. CNT toxicity within the Biomimetic interstitium chip was consistent with in vivo observations

Inhaled nanoparticles have been shown to pass through the epithelium barrier to the subepithelium and induce lung diseases by stimulating profibrotic responses and proinflammatory cytokine and chemokine secretion [60, 61]. Therefore, we explored the potential application of this lung interstitium chip for nanotoxicity studies. Of interest were MWCNTs because MWCNTs have been extensively assessed both *in vivo* and *in vitro* [20, 62, 63], which provided a benchmark for chip validation. The transport of MWCNTs through the epithelium and the lung inflammatory response were monitored after the epithelium was exposed to MWCNTs (5 μg/ml) for 24 h. The percentage of MWCNTs that crossed the epithelium into the interstitium on the chip was 2.24±1.80 %, significantly lower than the 7.61±2.85 % seen in the transwell model (Fig. 6a). This value is consistent with *in vivo* observations [20]. The culture medium was collected from the interstitium chamber and multiple inflammatory chemokines were measured. Although the expression of interleukin-4 (IL-4) and IL-6 was too weak to be detected, IL-8 expression associated with the fibroblasts was lower on the chip than the transwell model (Fig. 6b). The interstitium chip exhibited enhanced epithelial barrier function compared to the transwell models. As such, the penetration of MWCNTs across the epithelium was substantially reduced, and subsequently, the inflammatory responses of the fibroblasts to MWCNTs were alleviated, being in line with previous reports [20, 64, 65].

**Fig. 6.**
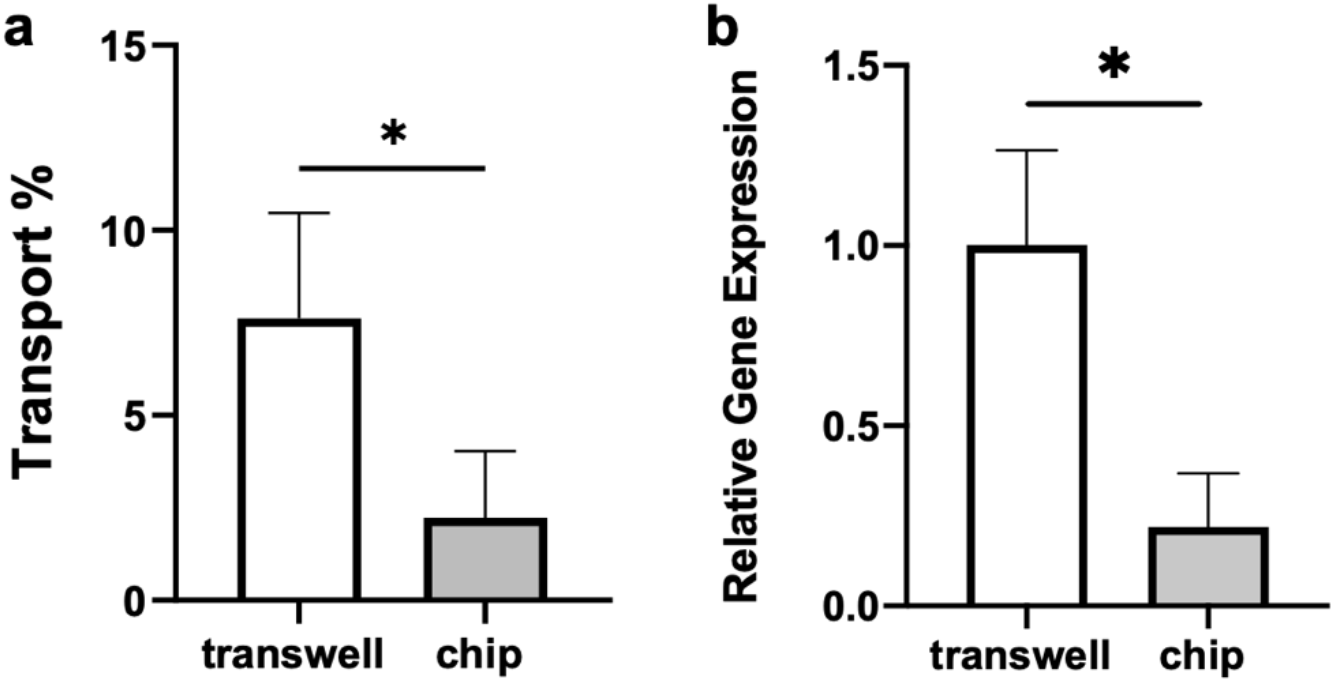
Toxicity assessment of MWCNTs on the chip and transwell model. (a) Percentage of MWCNTs penetrated across the epithelium. (b) Expression of IL-8 of the fibroblasts after the epithelium exposed to MWCNTs for 24 h, ^*^: *p* < 0.05 compared to the transwell model.

## 3. Conclusions

The lung alveolar interstitium chip described here closely imitated the key anatomical (epithelial cells co-cultured with fibroblasts encapsulated in 3D collagenous interstitial matrix via nanofibrous membrane) and physiological (interstitium matrix stiffness, interstitial flow, and 3D breathing-like mechanical stretch) characteristics of a human lung alveolus. The electrospun nanofibrous membrane promoted the formation of tight epithelium. Overall, the biomimetic interstitium chip demonstrated enhanced epithelial barrier function and extended longevity beyond 8 weeks with Col I-fibrin blend gels as the interstitium matrix. The key alveolar microenvironmental factors, such as nanofibrous membrane and interstitial matrix, can be finetuned to maintain the homeostasis of the epithelium and interstitium and thus further extend chip longevity. Importantly, the toxicity assessment of MWCNTs on the chip verified that the biomimetic interstitium chip represented a useful *in vitro* model for human lung alveolar interstitium. Looking forward, this interstitium chip system will have significant implications for drug development and disease modeling.

## 4. Materials and methods

### 4.1. Cell culture

Human alveolar epithelial cells (A549; Cat#: CCL-185, ATCC, Manassas, VA, USA) were cultured in Dulbecco’s Modified Eagle Medium (DMEM) with L-glutamine (Life Technologies, Carlsbad, CA, USA) supplemented with 10% fetal bovine serum (FBS; Sigma-Aldrich, St Louis, MO, USA), 100 U/ml penicillin and 100 μg/ml streptomycin (Life Technologies). Normal human lung fibroblasts (NHLFs; Cat#: CC-2512, Lonza, Walkersville, MD, USA) were cultured in fibroblast basal medium (Lonza) supplemented with FGM-2 SingleQuots supplements (Lonza), 100 U/ml penicillin, and 100 μg/ml streptomycin.

### 4.2. Fabrication and characterization of nanofibrous membranes

Nanofibrous membranes were fabricated by electrospinning a solution of PCL in 1,1,1,3,3,3-hexafluoro-2-propanol (HFIP, 10%, w/v). Briefly, the PCL solution was loaded in a syringe with a blunt-tipped needle as the spinneret and connected to a syringe pump. A 4” x 4” square glass plate was placed on a 3” x 3” aluminum-covered plate as a collector, which was placed 16 cm below the needle tip. The needle tip was connected to a 14 kV voltage and the aluminum-covered plate was grounded. The PCL solution was ejected at a flow rate of 0.5 ml/hour and the nanofibers were deposited on the collector.

The formed nanofibrous membranes were sputter-coated with gold using Denton Vacuum Desk V sputter coater (Denton Vacuum, Moorestown, NJ, USA) and imaged using a scanning electron microscope (SEM; TM3030 Plus, Hitachi High-Technologies Co., Tokyo, Japan). The fiber diameter and membrane pore size were analyzed using ImageJ (http://rsb.info.nih.gov/ij/index.html). For the fiber diameter, a line was drawn across the fiber, and then the length was measured. For the pore size, the images were adjusted using the threshold command and the area of each pore was analyzed using the analyze particles function. A histogram line graph was then generated to show the distribution of fiber diameters or pore sizes.

### 4.3. Fabrication of the lung interstitium chip

The chip consisted of an upper air chamber, a middle interstitium chamber, and a lower pneumatic chamber (Fig. 1), which had a concentric circle of 10 mm in diameter and different heights of 10, 2, and 5 mm, respectively. All the chambers were prepared by casting a mixture of PDMS resin and curing agent at a w/w ratio of 10:1.05 (Sylgard 184, Ellsworth Adhesives, Germantown, WA, USA) on their corresponding 3D printed mold and curing them at 75 °C for 2 h. To obtain the interstitium chamber with an open structure, the PDMS mixture was sandwiched between the 3D mold and a transparency film with a glass slide under a compressive pressure of approximately 1 MPa (illustrated in the red box in Fig. 1d). The pneumatic chamber was permanently bonded with a thin PDMS membrane via a μTA technique that we developed previously [40]. Briefly, the thin PDMS membrane was prepared by spin-coating a PDMS mixture on a silicon wafer at 2500 rpm for 1 min and curing it at 75 °C for 1 h. The pneumatic chamber with the opening facing down was then stamped on a thin PDMS layer, which was spin-coated on a silicon wafer at 4000 rpm for 30 s, and transferred onto the thin PDMS membrane, followed by curing at 75 °C for 1 h under a compressive pressure of approximately 1 MPa. Subsequently, the pneumatic chamber with the PDMS membrane was gently peeled off from the silicon wafer and was bonded with the middle interstitium chamber reversibly by applying the μTA technique again. For the reversible bonding, first, a 33% solution of PDMS mixture in hexane (Thermo Fisher Scientific, Waltham, MA, USA) was spin-coated on a silicon wafer at 4000 rpm for 2 min to form a thin adhesive layer, followed by prebaking at 50 °C for 5 min. Secondly, the interstitium chamber was stamped on the adhesive layer for 30 s and then transferred onto the pneumatic chamber, followed by curing at 75 °C for 1 h under a compressive pressure of approximately 1 MPa. Next, the nanofibrous membrane was reversibly bonded between the interstitium and air chambers following the aforementioned bonding process. One hole was punched in the pneumatic chamber and connected via the stainless-steel tubing to a programmable syringe pump (PHD 2000, Harvard Apparatus, Holliston, MA, USA), which provided cyclic mechanical stretch. Two holes were made in the air chamber as basal in and out ports for the culture medium perfusion through the interstitium chamber.

### 4.4. Characterization of mechanical strain

The mechanical strain (deformation) of the PDMS membrane on the pneumatic chamber with and without the interstitium layer was characterized using a previously described method [17]. Briefly, the deformation of the membrane or the membrane with interstitium layer was imaged using a charge-coupled device (CCD) camera (Model DMK 31; The Imaging Source, Charlotte, NC, USA). The captured images were adjusted using ImageJ to get a curvilinear profile, which was then digitalized using GetData Graph Digitalizer (http://www.getdata-graph-digitalizer.com). The measured linear strain was compared with the theoretical calculation as previously described [17].

### 4.5. Transwell culture models

Two transwell models were prepares for epithelial cell monoculture and co-culture as follows. Transwell monoculture models were prepared by attaching the nanofibrous membrane onto a transwell insert using the μTA technique described in *4.3*, and the epithelial cells were cultured on the apical side of the membrane. Transwell co-culture models were prepared as the control for the interstitium chip studies. NHLF at a seeding density of 2 × 10^5^ cells/cm^2^ were seeded onto the basal side of the transwell insert membrane (0.6 μm; Sterlitech, Auburn, WA, USA) and incubated for 1h. The insert was then inverted and A549 cells at 2 × 10^5^ cells/cm^2^ were placed in the upper compartment. For the ALI culture, the epithelial cell culture medium was removed after one week from the insert and the mixed epithelial and fibroblast culture media (1:1; v/v) was added to the low compartment and changed every three days.

### 4.6. Cell culture on lung interstitium chips

The chip was first oxygen plasma treated at the medium power for 1 min in a plasma cleaner (Model PDC-001; Harrick Plasma, Ithaca, NY, USA) and sterilized with 70% ethanol for 1 h, followed by rinsing with phosphate buffered saline (PBS, Sigma-Aldrich). Then, 500 μl of 50 μg/ml Col I solution (rat tail; Corning, Corning, NY, USA) was added to the cell culture chamber and interstitium chamber and incubated overnight in an incubator for the surface coating. The interstitium layer was prepared with Col I or Col I-fibrin blend hydrogel. In the case of Col I gels, Col I solution was prepared by mixing Col I, 10 × PBS, 1 N sodium hydroxide, and DMEM following the manufacturer’s protocol to obtain a final Col I concentration of 3 mg/ml. In the case of Col I-fibrin blend gels, fibrin solution was prepared by mixing solution A (a mixture of bovine fibrinogen (Sigma-Aldrich) and ɛ-aminocaproic acid (EACA, Sigma-Aldrich)) and solution B (a mixture of bovine thrombin (Sigma-Aldrich) and calcium chloride (CaCl_2_, Sigma-Aldrich)), with a final concentration of 5 mg/ml fibrinogen, 2 U/ml thrombin, and 5 mM CaCl_2_, and 2 mg/ml EACA. Subsequently, the fibrin solution was mixed with Col I solution (3 mg/ml) and human lung fibroblasts (at a seeding density of 2.0 × 10^5^ cells/mL), and the mixture was injected into the interstitium chamber and incubated at 37 °C for 4 h to form the cell-laden hydrogel matrix. The A549 cells at a density of 2.0 × 10^5^ cells/cm^2^ were then added into the air chamber and cultured under static condition for three days before the interstitial flow and cyclic mechanical stretch were applied. After cultured for one week under the mechanical stretch, the epithelial cell culture medium was aspirated from the air chamber and a mixed epithelial and fibroblast culture media (1:1, v/v) supplemented with EACA (2 mg/ml) was perfused through the interstitium chamber.

### 4.7. Immunofluorescent staining

The cells were fixed with 1:1 (v/v) of methanol: acetone at −20 °C for 10 min and blocked with tris buffered saline (TBS) (KD Medical, Columbia, MD, USA) containing 10% (v/v) FBS, 10% (v/v) goat serum (Sigma-Aldrich), and 1% bovine serum albumin (BSA; Akron Biotech, Boca Raton, FL, USA) at room temperature for 30 min. After blocking, the cells were incubated with primary antibody at 4 °C overnight followed by secondary antibody incubation at room temperature for 1 h. The information of antibodies used for immunostaining and western blot were listed in Table S1. The nuclei were stained with SlowFade™ Gold Antifade Mountant with DAPI (Life Technologies) and the samples were examined using a Nikon Ti eclipse fluorescence microscope (Nikon, Melville, NY, USA).

### 4.8. Western blotting

The total proteins were extracted by lysing the cells with radioimmune precipitation assay (RIPA) buffer (Santa Cruz Biotechnology, Santa Cruz, CA, USA) that contained protease inhibitor for 30 min on ice followed by centrifuging at 4 °C, 12,000 rpm for 5 minutes and collecting the supernatant. Proteins were separated by 8% sodium dodecyl sulfate-polyacrylamide gel electrophoresis (SDS-PAGE) and transferred onto polyvinylidene fluoride (PVDF) membrane (Thermo Fisher Scientific). The membranes were blocked in 5% nonfat milk (RPI International, Mount Prospect, IL, USA) dissolved in TBS with 0.1% Tween-20 (Thermo Fisher Scientific) at room temperature for 1 h. Subsequently, the membrane was incubated with primary antibody and horseradish peroxidase (HRP) – conjugated secondary antibody. Protein bands were visualized by ECL detection kit (EMD Millipore, Burlington, MA, USA) and images were acquired using the ChemiDoc MP image system (Bio-Rad, Hercules, CA, USA). The band intensity was quantified using ImageJ and normalized to the expression of β-actin.

### 4.9. Permeability assay

The epithelial permeability was measured in the 12-well transwell plate-based models including the epithelial cell monoculture on a nanofibrous membrane and co-culture with fibroblast cells and on the interstitium chips. In the transwell models, 500 μl of 4 kDa (100 μg/ml) or 70 kDa (500 μg/ml) fluorescein isothiocyanate (FITC)-dextran (Sigma-Aldrich) was added to the insert and 1 ml of cell culture medium was added to the low compartment. The fluorescence intensity of the medium from the insert and the low compartment was measured after incubation for 2 h. On the chip, the interstitium perfusion was halted and 200 μl of FITC-dextran was added into the air chamber. The chip was incubated for 2 h in an incubator and then the gel and medium in the interstitium were collected after the chip was dissembled for fluorescence intensity measurements.

The concentration of FITC-dextran was determined by fluorescence intensity that was compared to a calibration curve of FITC-dextran concentration vs fluorescence intensity. The apparent permeability *P_app_* was calculated as 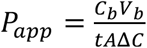, where *C_b_* and *V_b_* are the dextran concentration and solution volume in the low compartment of the transwell or interstitium chamber of the chip, respectively, *t* is the diffusion time, *A* is the total area of diffusion, *ΔC* is the concentration change across the barrier [66]. The permeability coefficient of the epithelial barrier, P_epi_, was determined from the *P_app_* and the background permeability coefficient *P_o_* (measured in a transwell plate or chip without epithelium) as follows: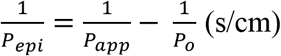.

### 4.10. SEM imaging

The epithelial cells were fixed *in situ* with 4% paraformaldehyde (PFA, Sigma-Aldrich) and 2% glutaraldehyde solution (Fisher Chemical, Fairlawn, NJ, USA) at room temperature for 4 h and then dehydrated with gradient ethanol (30%, 50%, 70%, 80%, 90%, 95%, and 100%) and then hexamethyldisilazane (HMDS), each step being 10 min. The epithelial cell layer on the nanofibrous membrane or the transwell membrane was taken off, sputter-coated by gold, and imaged as detailed in *4.2*.

### 4.11. CNT penetration assay

MWCNTs (XNRI MWNT-7; Mitsui & Company, NY, USA) were dispersed in PBS containing 5 mg/ml of BSA and diluted in culture medium to a concentration of 5 μg/ml as previously reported [27]. The MWCNT suspension was added to the transwell insert or the chip air chamber, similar to the permeability assay in *4.9* with the MWCNT suspension replacing the FITC-dextran. After incubation at 37 °C for 24 h, the medium in the transwell low compartment was collected for optical density (OD) measurement. The gel and medium in the interstitium chamber of the chip were collected and digested with 4 mg/ml collagenase in PBS at 37 °C before the OD measurement. The OD 640 value of the solution was measured and MWCNT concentration was calculated by normalizing the OD value with the value obtained from the standard curve and thus the relative transport across the barrier was calculated.

### 4.12. Real-time quantitative reverse transcription-polymerase chain reaction (qRT-PCR) assay

After MWCNT treatment, the expression of transcripts for the pro-inflammatory cytokines IL-4, 6, 8 derived from the NHLFs on the chip and transwell cultures were analyzed by qRT-PCR assay. The NHLF in the interstitium of the chip was collected by incubating with 4 mg/ml collagenase at 37 °C and centrifuged for 5 min at 400 *g* at 4 °C. Total RNA was extracted using the Aurum Total RNA Mini Kit (Bio-Rad) and cDNA was synthesized using iScript RT Supermix (Bio-Rad). The qRT-PCR samples were prepared using SsoAdvanced Univ SYBR Grn Suprmix (Bio-Rad). The reaction was performed using a CFX96 Touch Real-Time PCR Detection System (Bio-Rad) under the following conditions: 40 cycles of 95 °C for 15 s, 60 °C for 30 s, followed by a melt curve of 65-95 °C at 0.5 °C increments, at 2-5 sec/step. The primer sequences used are in Table S2. mRNA expression was normalized to α-tubulin mRNA expression.

### 4.13. Statistical analysis

The data were presented as mean ±standard error of the mean (S.E.M.). The statistical differences were analyzed using two-tailed *t* test using Prism 8 (GraphPad software, San Diego, CA, USA). Statistically significant differences were considered at a level of *p* < 0.05.

## Credit author statement

KM, JL, and CL: Investigation, methodology, analysis, and visualization. KM, CC, and YY: Data interpretation. KM, MS, BM, and YY: Writing. YY: Conceptualization, methodology, and supervision.

## Data availability

Data will be made available on request.

## Acknowledgments

This work is partly supported by National Institutes of Health R15GM122953 (to Y.Y.).

## Appendix A. Supplementary data

Supplementary data to this article can be found online at xxx

