## Supplemental Materials for "Biomimetic human lung alveolar interstitium chip with extended longevity"

**Computational simulation of interstitial velocity and shear stress**

The design of the interstitium chamber and computational fluid dynamics simulation of the interstitial velocity and shear stress were performed using SOLIDWORKS 2021 (Dassault Systems SOLIDWORKS Co., Waltham, MA, USA). The chamber design and dimensions were shown in Fig. S1a, b. For an interstitial perfusion volume flow rate (*i.e.*, 20 μl/hr in this study), the fluidic velocity and shear stress were simulated and exported using the built-in Flow Simulation module. For the simulation, the porous media function was applied to the chamber to mimic the collagen matrix. The porosity, pore size setting and other parameter inputs are given in the table below. In the Simulation module, a 95% porosity was used to estimate the porous structure of collagen, and thus the simulation results were the approximation. For comparison, additional setting where the chamber was filled with water was also simulated. The comparisons of interstitial velocity and shear stress profiles are shown in Fig. S1c-e.

| **Input settings** | **Value** |
|---|---|
| Porosity of collagen | 95% |
| Pore size | 10um |
| Gravity | 9.81m/s$^2$ |
| Temperature | 310.2K |
| Density of Water | 998.1kg/m$^3$ |
| Dynamic viscosity | 0.00066238Pa*s |

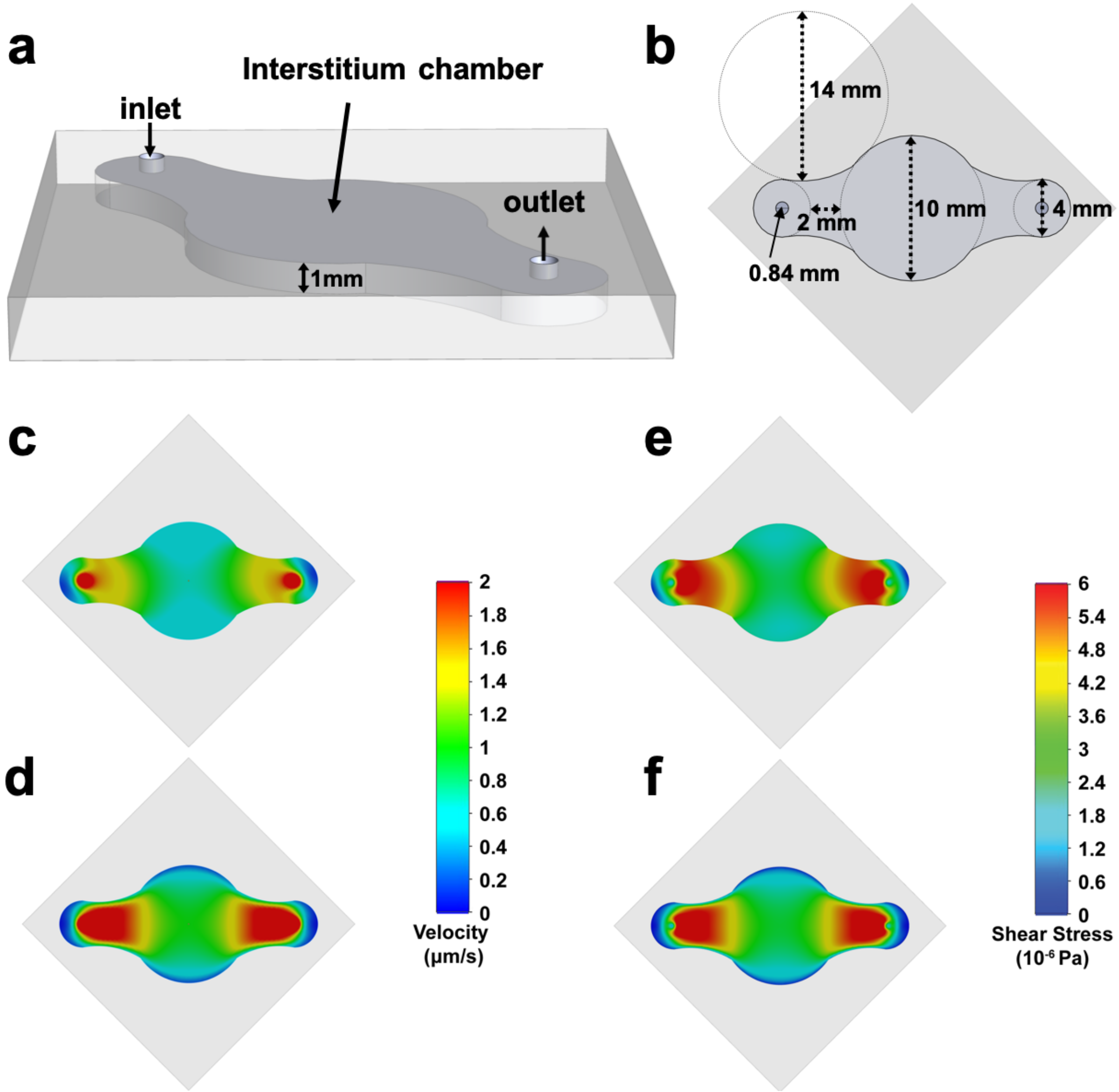


**Figure S1.** Solidworks simulation of the interstitial fluidic flow. (a) Perspective and (b) top views of the interstitium chamber. (c, d) interstitial fluidic velocity and (e, f) shear stress profiles of interstitium chamber filled (c, e) with the collagen gel or (d, f) water at a constant interstitial volumetric flow rate of 20 μl/hr.

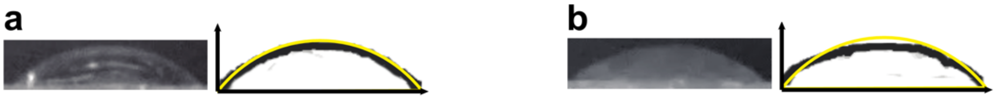


**Figure S2.** Characterization of mechanical stretches. Original CCD images (left) and the profiles (black curve in the right) of the deformed (a) PDMS membrane bonded on the pneumatic chamber and (b) nanofibrous membrane on the interstitial layer bonded on the pneumatic chamber under a theoretical strain of 15% (yellow curve).

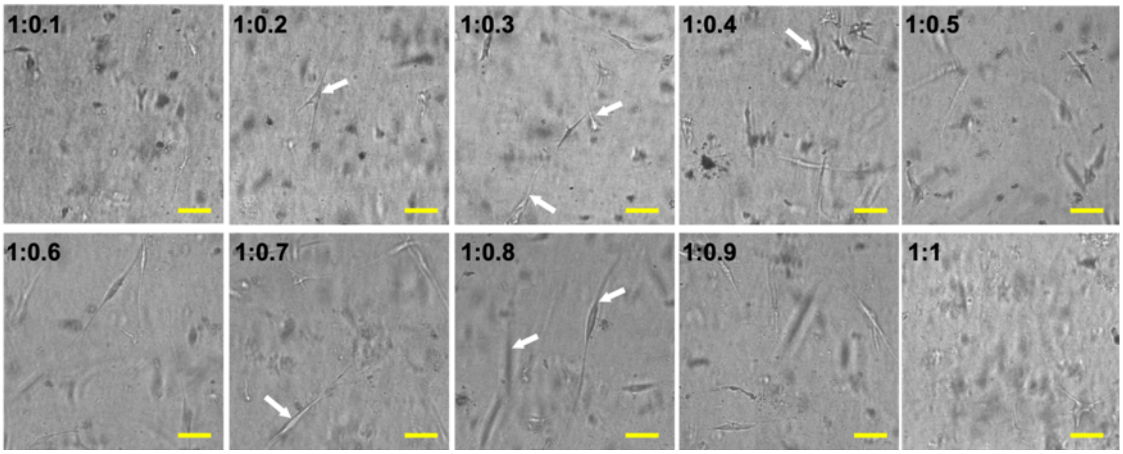


**Figure S3.** NHLF cultured in Col I-fibrin blend gels for 14 days with Col I: fibrin ratio ranging from 1:0.1 to 1:1. Scale bars: 100 μm. The white arrows indicated the cells.

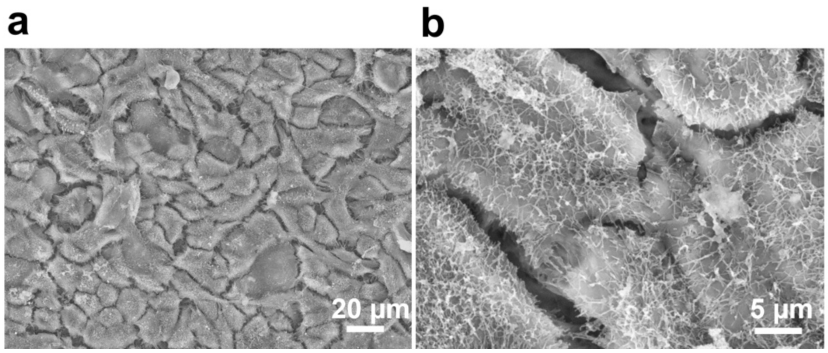


**Figure S4.** SEM images of epithelial cell layer in the transwell model with (a) low and (b) high magnification after 8-week culture.

**Supporting videos:**

**Video S1** Deformation of PDMS membrane bonded on the pneumatic chamber under a theoretical 15% strain.

**Video S2** Deformation of nanofibrous membrane on the interstitial layer bonded on the pneumatic chamber under a theoretical 15% strain.

**Supporting tables:**

**Table S1** Antibodies for western blot and immunofluorescence staining

| Antibody | Vendor | Catalog # | Dilution |
|---|---|---|---|
| ZO-1 | Life Technologies | 339100 | 1:100 (IF)<br>1:500 (WB) |
| ZO-3 | Life Technologies | 364000 | 1:200 (WB) |
| E-cadherin | Life Technologies | 131700 | 1:1000 (WB) |
| Occludin | Life Technologies | 331594 | 1:1000 (WB) |
| β-actin | Life Technologies | AM4302 | 1:1000 (WB) |

**Table S2** Primers for qRT-PCR

| Gene | Primer sequence |
|---|---|
| IL-4 | Forward: TCTTTGCTGCCTCCAAGAACA<br>Reverse: GTAGAACTGCCGGAGCACAG |
| IL-6 | Forward: TCCGGGAACGAAAGAGAAGC<br>Reverse: GAGAAGGCAACTGGACCGAA |
| IL-8 | Forward: ACACTGCGCCAACACAGAAA<br>Reverse: CAACCCTCTGCACCCAGTTT |
| TUBA1A | Forward: CGGGCAGTGTTTGTAGACTTGG<br>Reverse: CTCCTTGCCAATGGTGTAGTGC |
